# Elevated rates and biased spectra of mutations in anaerobically cultured lactic acid bacteria

**DOI:** 10.1101/2025.02.28.639667

**Authors:** Owen F. Hale, Michelle Yin, Megan G. Behringer

## Abstract

The rate, spectrum, and biases of mutations represent a fundamental force shaping biological evolution. Convention often attributes oxidative DNA damage as a major driver of spontaneous mutations. Yet, despite the contribution of oxygen to mutagenesis and the ecological, industrial, and biomedical importance of anaerobic organisms, relatively little is known about the mutation rates and spectra of anaerobic species. Here, we present the rates and spectra of spontaneous mutations assessed anaerobically over 1000 generations for three fermentative lactic acid bacteria species with varying levels of aerotolerance: Lactobacillus acidophilus, Lactobacillus crispatus, and Lactococcus lactis. Our findings reveal highly elevated mutation rates compared to the average rates observed in aerobically respiring bacteria with mutations strongly biased towards transitions, emphasizing the prevalence of spontaneous deamination in these anaerobic species and highlighting the inherent fragility of purines even under conditions that minimize oxidative stress. Beyond these overarching patterns, we identify several novel mutation dynamics: positional mutation bias around the origin of replication in Lb. acidophilus, a significant disparity between observed and equilibrium GC content in Lc. lactis, and repeated independent deletions of spacer sequences from within the CRISPR locus in Lb. crispatus providing mechanistic insights into the evolution of bacterial adaptive immunity. Overall, our study provides new insights into the mutational landscape of anaerobes, revealing how non-oxygenic factors shape mutation rates and influence genome evolution.

## Introduction

Life on Earth began in an anaerobic environment that remained so for approximately 2 billion years (Lyons et al. 2014). With the advent of oxygenic photosynthesis, many organisms adapted to harness the redox potential of molecular oxygen to more efficiently generate energy from available nutrients (Raymond and Segrè 2006; Soo et al. 2017). However, aerobic respiration evolved on the preexisting chassis of anaerobic respiration, leaving anaerobes vulnerable to oxidative stress caused by and targeted to the highly reactive enzymes needed for efficient anaerobic metabolism (Khademian and Imlay 2021; Glass et al. 2023). In response, some species evolved behaviors and/or complex machinery for protecting their cells from oxidative damage, while others remodeled their physiology to tolerate transient exposure to oxygen (Khademian and Imlay 2021). In addition to the historical importance of anaerobes and the transition to an aerobic atmosphere, anaerobes today represent important environmental (Tiedje et al. 1984; Op den Camp et al. 2006), industrial (Hatti-Kaul and Mattiasson 2016), and pathogenic (Church 2016) members of the biosphere, and the role of oxygen in driving evolution through mutagenesis has been well characterized (Cooke et al. 2003; Foster et al. 2015). Despite this, little is known about the mutation rates and spectra of anaerobic organisms.

Characterizing the rate at which mutations occur and the relative rates of specific subtypes of mutations, also known as the mutation spectrum, is critical for biological research as mutation provides the raw material of genetic diversity on which the other forces of evolution can act, is directly influenced by a species’ evolutionary past, and constrains a species’ evolutionary future. To date, mutational studies have been completed for over 150 organisms across the tree of life and scale of organismal complexity, providing a deep understanding of mutation rates and highlighting the role of oxidative damage in shaping the mutation rate and spectrum (Lynch et al. 2023). However, despite the curation of such a rich dataset of mutational patterns, quantification of mutation rate and spectrum has primarily focused on species with aerobic metabolisms under oxygenated conditions. Because of the requirement for specialized equipment and the fastidious and/or pathogenic nature of many anaerobes, mutation rates have only been directly quantified for two anaerobically cultured organisms to date (Shewaramani et al. 2017; Gu et al. 2021) As anaerobes include important environmental, industrial, and pathogenic taxa and occupy a wide range of phylogenetic and ecological diversity, there exists a need to thoroughly investigate the causes and consequences of mutation across the anaerobic tree of life.

Lactic acid bacteria (LAB) are a well-suited group of organisms with which to begin this work. LAB are a polyphyletic group of bacteria known for their production of lactic acid and include, among others, members of the order Lactobacillales (Wang et al. 2021). Most LAB are classified as aerotolerant or facultative anaerobes with fermentative lifestyles and can be found in diverse habitats from plants to insects to mammals (Kleerebezem et al. 2020; Zheng et al. 2020). Importantly, many have been linked to beneficial health effects in humans (Canani et al. 2007; Lebeer et al. 2008; Stapleton et al. 2011; Smith and Ravel 2017) and are used in industrial processes that span dairy fermentation (Song et al. 2017; Stoyanova et al. 2023) and the production of recombinant therapeutics (Song et al. 2017). The Lactobacillales are generally incapable of performing aerobic respiration, and those that can often cannot construct a complete electron transport chain without exogenous heme (Duwat et al. 2001; Zotta et al. 2017). These organisms also vary in their susceptibility to oxidative stress and possess few genes encoding enzymes that mitigate oxidative damage, such as catalase (**Fig. 1**). The diversity of these organisms, both in their lifestyles and physiologies, as well as their low frequency of pathogenicity and ease of culture, makes them an ideal system for studying the evolution of the mutation rate and spectrum.

**Figure 1.**
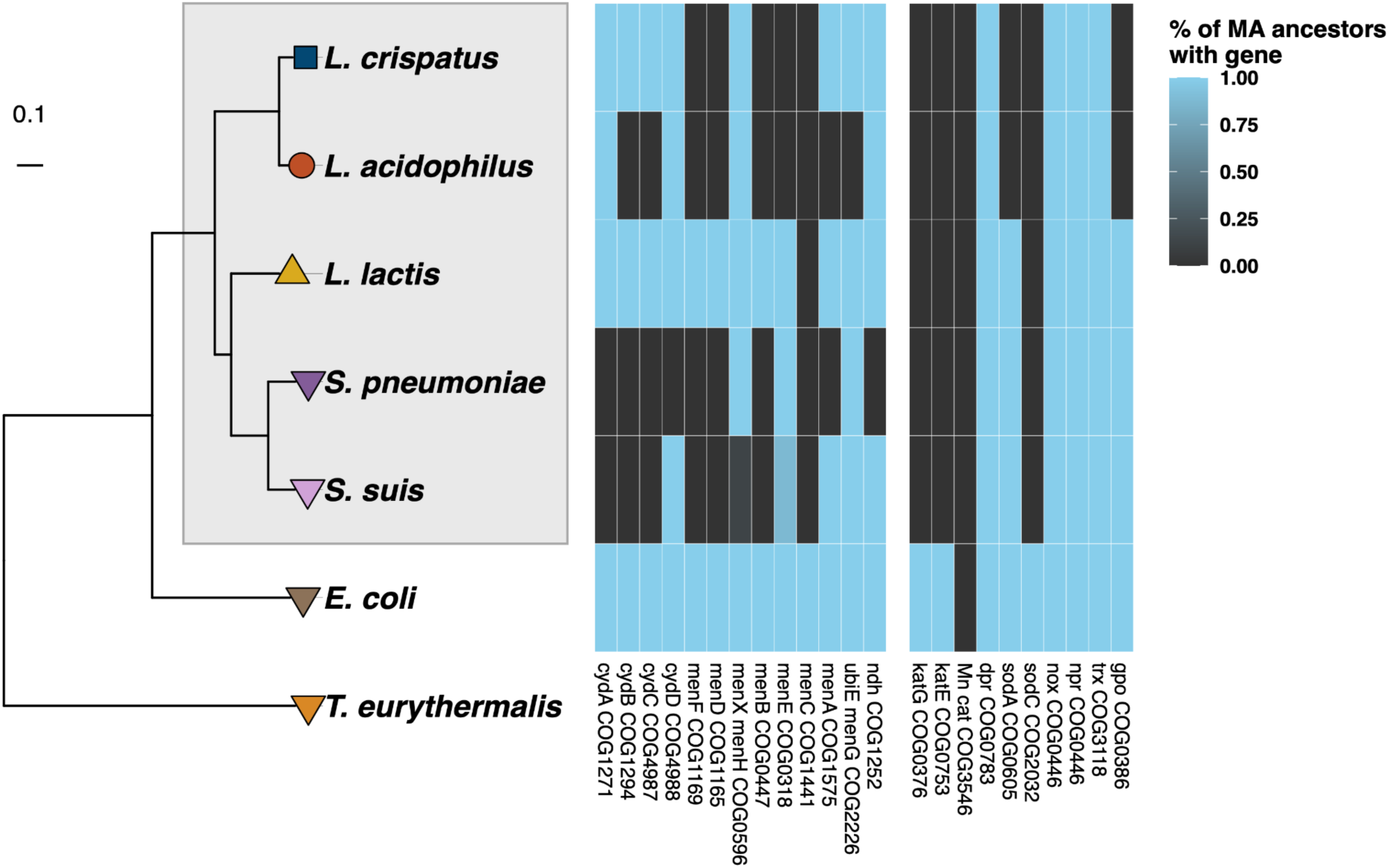
Phylogenetic relatedness and oxidative stress-related gene carriage of notable prokaryotes with mutation rates measured via MA/WGS. Maximum likelihood phylogeny of prokaryotes with mutations measured in this (*Lb. acidophilus*, *Lb. crispatus*, and *Lc. lactis*) and previous (*E. coli*, *S. pneumoniae*, *S. suis*, and *T. eurythermalis*) MA/WGS experiments constructed from the *rpoB* gene. LAB are highlighted in the gray box. The heatmap shows the fraction of ancestral genomes used in each MA/WGS experiment that contain genes involved in aerobic respiration (left gene set) and scavenging reactive oxygen species (right gene set). For *S. suis*, n = 8, while n = 1 for all other taxa. *T. eurythermalis* was excluded from the heatmap due to its evolutionary divergence from bacteria.

Mutation accumulation experiments coupled with whole-genome sequencing (MA/WGS), the gold standard method for measuring an organism’s mutation rate and spectrum, have been performed on three aerobically cultured LAB: *Lactococcus lactis* (Long et al. 2018), *Streptococcus pneumoniae* (Jiang et al. 2024), and *Streptococcus suis* (Murray et al. 2021). In all three instances, high mutation rates and biased mutation spectra were observed, although it is unclear whether these observations are the result of susceptibility to oxidative DNA damage rather than an intrinsically higher mutation rate. In MA/WGS experiments, replicate populations of isogenic microbial strains are continuously cultured through single-cell bottlenecks, whole-genome sequenced, and compared to the experiment’s ancestral strain to identify new mutations (Mukai 1964; Halligan and Keightley 2009). This allows for the accumulation of spontaneous mutations and minimizes the influence of selection on the observed spectrum of mutations. MA/WGS confers advantages relative to other methods such as fluctuation tests because it reliably and repeatably measures the mutation rate and spectrum across the entire genome, enabling the observation of genome-wide mutational patterns (Long et al. 2015; Dettman et al. 2016; Behringer and Hall 2016; Dillon et al. 2018; Kucukyildirim et al. 2020). The wealth of data generated by MA/WGS experiments has enabled evolutionary theorists to test hypotheses about the determinants of the mutation rate. The current leading hypothesis concerning the evolutionary factors shaping mutation rates is the drift barrier hypothesis, which predicts that selection will act to minimize mutation rates with the power of genetic drift limiting the efficacy of selection in small populations (Lynch 2010). Given that many LAB are adapted to living within a host (Duar et al. 2017), a lifestyle associated with a decreased effective population size (N_e_) and therefore increased drift (Bobay and Ochman 2018), we expect host-adapted LAB to have high mutation rates relative to free-living bacteria.

Here, we performed MA/WGS experiments on three anaerobically cultured members of the order Lactobacillales: *Lactobacillus acidophilus*, a vertebrate gut commensal (Gilliland et al. 1975; Bull et al. 2013) used in dairy fermentations (Sanders and Klaenhammer 2001; Bull et al. 2013); *Lactobacillus crispatus*, a commensal of the human gut (Pan, et al. 2020a; Pan, et al. 2020b) and urogenital tract (Smith and Ravel 2017; Pan, et al. 2020a; Pan, et al. 2020b; Song et al. 2022) as well as the guts of poultry (Edelman et al. 2002; Pan, et al. 2020a) and sometimes found in sourdough starter (Li et al. 2020); and *Lactococcus lactis* subsp. *lactis*, a human gut commensal used in dairy and other industrial fermentation processes (Long et al. 2018; Kleerebezem et al. 2020). For all three species, we calculated the per base pair substitution rate, per base pair indel rate, and genome-wide mutational biases and characterized the mutation spectra and occurrence of structural variants. We compare these results to aerobically cultured *Escherichia coli* (Lee et al. 2012), three previously studied but aerobically cultured Lactobacillales (Long et al. 2018; Murray et al. 2021; Jiang et al. 2024), and the anaerobically cultured hyperthermophilic archaeon, *Thermococcus eurythermalis* (Gu et al. 2021). To our knowledge this work represents the most extensive characterization of anaerobic mutation rates to date, markedly increasing our understanding of how the presence or absence of oxygen influences the evolutionary and mutational landscape.

## Results

### Anaerobically cultured LAB exhibit extremely high SNM rates

Mutation accumulation for 66 lines of *Lb. acidophilus*, 66 lines of *Lb. crispatus*, and 50 lines of *Lc. lactis* was conducted over 1,171, 1,084, and 1,067 generations, respectively. Across these lines, we observed a total of 3,009 single nucleotide mutations (SNM) in *Lb. acidophilus*, 140 in *Lb. crispatus*, and 107 in *Lc. lactis*, for a genome-wide per base pair per generation substitution rate of 1.97×10^−8^ (Standard error (SE): 9.79×10^−11^), 9.01×10^−10^ (SE: 1.11×10^−11^), and 8.36×10^−10^ (SE: 1.24×10^−11^), respectively (**Table 1**, **Fig. 2A**). These rates are considerably higher than previously examined bacterial SNM rates such as the *E. coli* K-12 rate (Lee et al. 2012) of 2.2×10^−10^, but similar to the SNM rates of *Lc. lactis* (Long et al. 2018), *S. pneumoniae* (Jiang et al. 2024), and *S. suis* (Murray et al. 2021) in aerobic conditions, suggesting that these higher mutation rates may be a general feature of LAB (**Fig. 2C**).

**Figure 2.**
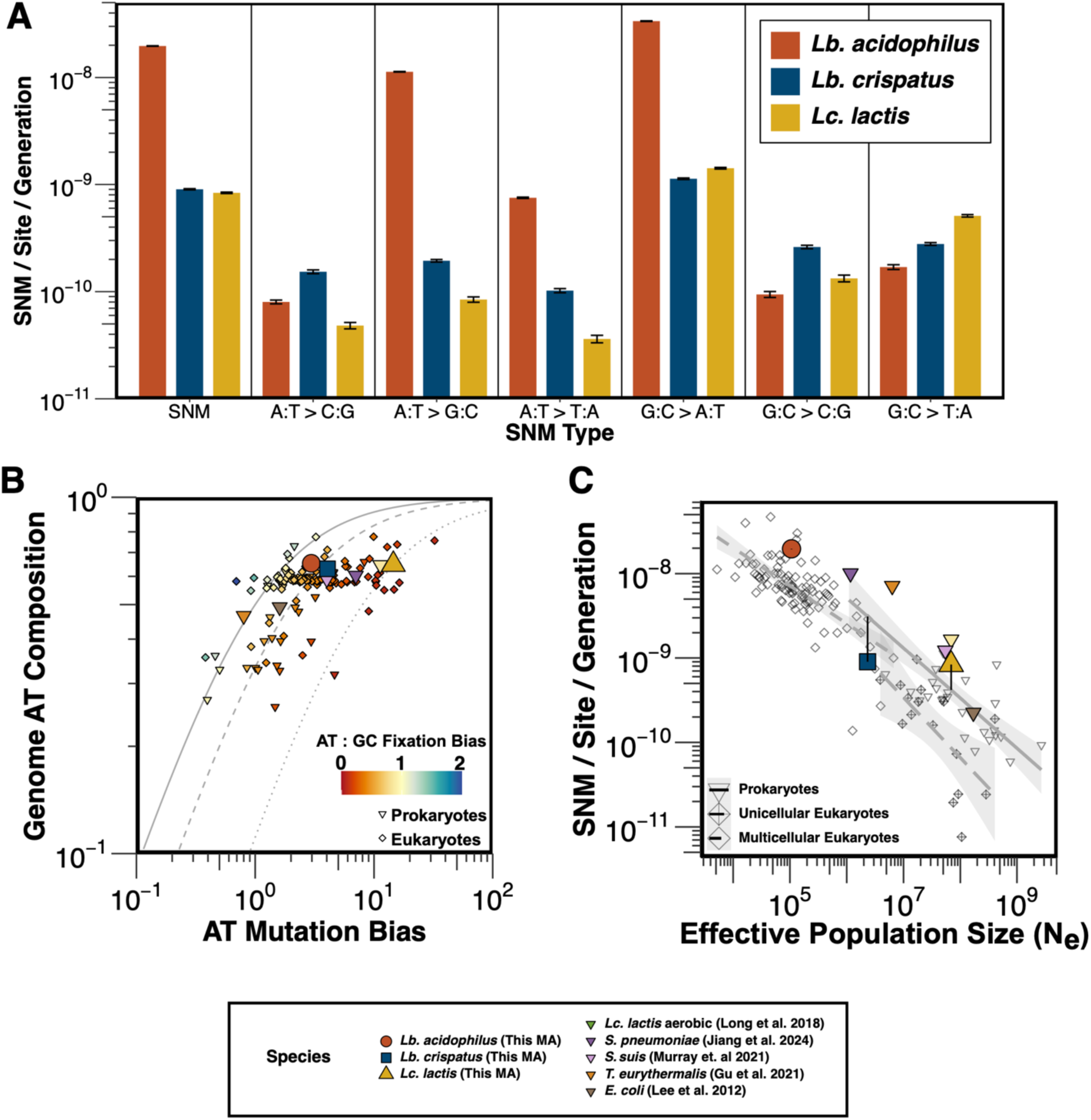
High rates and biased spectra of SNMs in anaerobically cultured LAB. (A) Per site per generation rates of all SNMs and each of the 6 SNM types. (B) Relationship between AT mutation bias, AT composition, and AT fixation bias in diverse organisms. Reference lines for AT:GC fixation biases of 1:1 (solid), 1:2 (dashed), and 1:8 (dotted) are included. (C) Relationship between N_e_ and mutation rate, which is consistent with the drift barrier hypothesis of mutation rate evolution. Organisms are split into prokaryotes (solid line and inverted triangles), unicellular eukaryotes (long dashed line and crossed diamonds), and multicellular eukaryotes (short dashed line, empty diamonds). For *Lb. acidophilus*, *Lb. crispatus*, and *Lc. lactis*, the difference between the observed mutation rate and the mutation rate predicted by the linear regression model created with previously reported prokaryotic mutation rates and Ne values is represented by a black line. For a description of the external data used in panels B and C see the Materials and Methods.

**Table 1.**
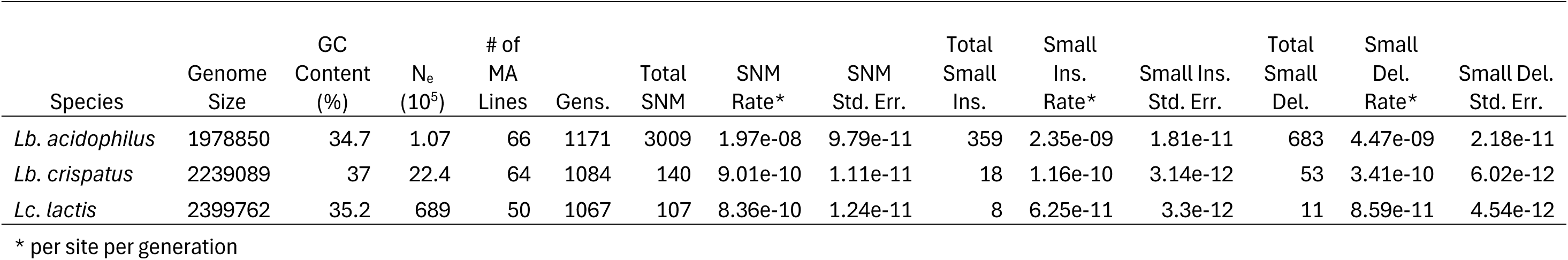
Genomic information, experimental parameters, and observed mutation counts and rates.

Minimizing the effects of selection on mutations is essential for MA/WGS experiments to accurately reflect an organism’s spontaneous mutation rate and spectrum in a given environment. We accomplished this by repeatedly bottlenecking each MA line to reduce the N_e_, which we calculated as the harmonic mean of the number of cells at each generation between bottlenecks, assuming non-overlapping generations, a bottleneck size of one cell, and that every cell in every generation divides into two individuals. We determined that the N_e_ during MA was 11, 11, and 12 for *Lb. acidophilus, Lb. crispatus,* and *Lc. lactis*, respectively. As the minimum selection coefficient *s* required to overcome random drift is 1/N_e_ in haploid organisms (Lynch et al. 2016), new mutations would need an extremely high absolute *s* of at least 0.09 to bias our results.

We performed multiple tests to determine if selection significantly biased the set of observed mutations during mutation accumulation. First, we tested whether the observed SNM, INS, and DEL counts in each species fit a negative binomial distribution, a generalization of the Poisson distribution that allows for greater variances. We found no evidence of deviation from the expected distribution (*χ*^2^ all p > 0.05) (**Supplemental Fig. S1**). Second, we confirmed that SNM mutations within coding sequences of all three species did not occur more frequently than would be expected by chance (*χ*^2^ all p > 0.05).

### SNM spectra are heavily biased towards G:C → A:T

We observed high SNM rates in all three LAB species relative to previous MA/WGS experiments in other bacterial species, primarily driven by G:C → A:T transitions and suggesting oxidative damage as a potential cause of increased SNM rates (**Fig. 2A**). However, each species varied in the degree to which their mutation spectra were biased toward G:C → A:T transitions. In *Lb. acidophilus*, the rate of A:T → G:C transitions was only slightly lower than G:C → A:T transitions, resulting in an extremely high ratio of transition to transversion mutations (Ts/Tv) of 30.02 but a relatively low AT mutation bias (the ratio of the rate of SNMs converting GC sites to AT sites to the rate of SNMs converting AT sites to GC sites) of 2.97 (**Fig. 2B**). In contrast, the mutation spectrum of *Lb. crispatus* is much more balanced as the rate of G:C → A:T mutations was only 11 times greater than its least frequent SNM. Thus, *Lb. crispatus* exhibited a Ts/Tv ratio of 1.50 and an AT mutation bias of 4.01, values well within the normal range for bacteria. Interestingly, *Lc. lactis* exhibits a relatively high frequency of G:C → T:A mutations and G:C → A:T mutations, leading to a greater AT mutation bias of 14.53 than would be expected given its genomic GC-content of 35.24% and indicating there is a high fixation bias towards new GC alleles over AT alleles of 7.91 in *Lc. lactis*.

Additionally, we can compare the SNM rate and spectrum we observed in anaerobically cultured *Lc. lactis* to those observed in a prior MA/WGS study of *Lc. lactis* cultured in aerobic conditions (Long et al. 2018). The largest changes in the mutation spectrum between the two experiments are in SNMs at G:C sites, where the observed mutation rates are much lower in anaerobic than aerobic conditions. As these sites are particularly susceptible to mutation due to oxidative damage through cytosine deamination (Nabel et al. 2012) and 8-oxoguanine formation (Grollman and Moriya 1993), these results are consistent with the hypothesis that the biased mutation spectrum of LAB is due to oxidative damage.

### SNM rates in LAB are consistent with the drift barrier hypothesis of mutation rate evolution

To determine if mutation rates of anaerobically cultured LAB conform to the predictions of the drift barrier hypothesis of mutation rate evolution, the same scaling law observed of aerobic species, we investigated the relationships between the mutation rates observed in this study with estimates of each species’ N_e_. We expect the N_e_ of the LAB species in this study to be relatively small due to their domestication for industrial use and host-associated lifestyles (Moyers et al. 2018; Bobay and Ochman 2018). We can predict N_e_ for *Lb. acidophilus* and *Lb. crispatus* using nucleotide diversity estimates from multilocus sequence typing (MLST) of *Lactobacillus delbrueckii* (Song et al. 2016), the type species of the genus *Lactobacillus*, which has a similar lifestyle to *Lb. acidophilus* and *Lb. crispatus* (Song et al. 2016; Ghosh et al. 2020; Vaughan et al. 2021). Assuming the relationship for haploid organisms, N_e_ = π/2μ where π is pairwise diversity and μ is the per site per generation mutation rate (Tajima 1983), we estimate the N_e_ for *Lb. acidophilus* and *Lb. crispatus* as 1.07 x 10^5^ and 2.33 x 10^6^, respectively. These calculated N_e_ for *Lb. acidophilus* and *Lb. crispatus* are relatively small compared to the aerobic bacterial species previously evaluated with MA/WGS averaging ∼ 1 x 10^8^ (Lynch et al. 2023). For example, *Escherichia coli*, a species with a greater aerobic capacity and aerotolerance characteristic of its biphasic host / environment-associated lifestyle, has an estimated N_e_ ranging from 1.5 - 4.5 x 10^8^, which is ∼10^2^ and ∼10^3^ times larger than *Lb. acidophilus* and *Lb. crispatus*, respectively (Berg 1996; Sung et al. 2012; Sung et al. 2016; Bobay and Ochman 2018; Moyers et al. 2018).

Alternatively, for *Lc. lactis*, we expect a slightly greater N_e_ than in *Lb. acidophilus* or *Lb. crispatus* due to *Lc. lactis*’s more free-living lifestyle on plants and soils in addition to its domesticated strains (Urbach et al. 1997; Nomura et al. 2006). MLST studies and estimates of N_e_ predict a N_e_ of 6.89 x 10^7^ and confirm this expectation(Passerini et al. 2010; Xu et al. 2014; Liu et al. 2022; Lynch et al. 2023). Thus, while the mutation rates of anaerobically cultured LAB initially appear unexpectedly high, given these estimates of N_e_, our results are consistent with the drift barrier hypothesis of mutation rate evolution (Sung et al. 2012) (**Fig. 2C**).

### Lb. acidophilus G:C → A:T mutation rates are elevated near the replication origin

Previous MA/WGS experiments conducted with mutator strains have identified variations in the mutation rate across the length of the bacterial chromosome in which SNM rates peak approximately halfway between the origin of replication (*oriC*) and the replication terminus (Long et al. 2015; Dettman et al. 2016; Dillon et al. 2018). To effectively detect locational biases in mutation rates, mutator strains are typically required to generate a sufficient number of mutations to provide the analysis enough power to achieve statistical significance. Thus, the high SNM rate of *Lb. acidophilus* allowed us to screen for locational biases, revealing a 3x increase in G:C → A:T transitions close to the *oriC* relative to the terminus (*ρ* = −0.53, Bonferroni corrected p = 1.36 ×10^−15^) (**Figs. 3A & 3B**). Interestingly, this pattern is not even qualitatively present in the rest of the mutation spectra of *Lb. acidophilus* nor in the mutation spectra of *Lb. crispatus* and *Lc. lactis* (**Supplementary Fig. S1**, all |*ρ*| < 0.15, all Bonferroni corrected p > 0.1).

**Figure 3.**
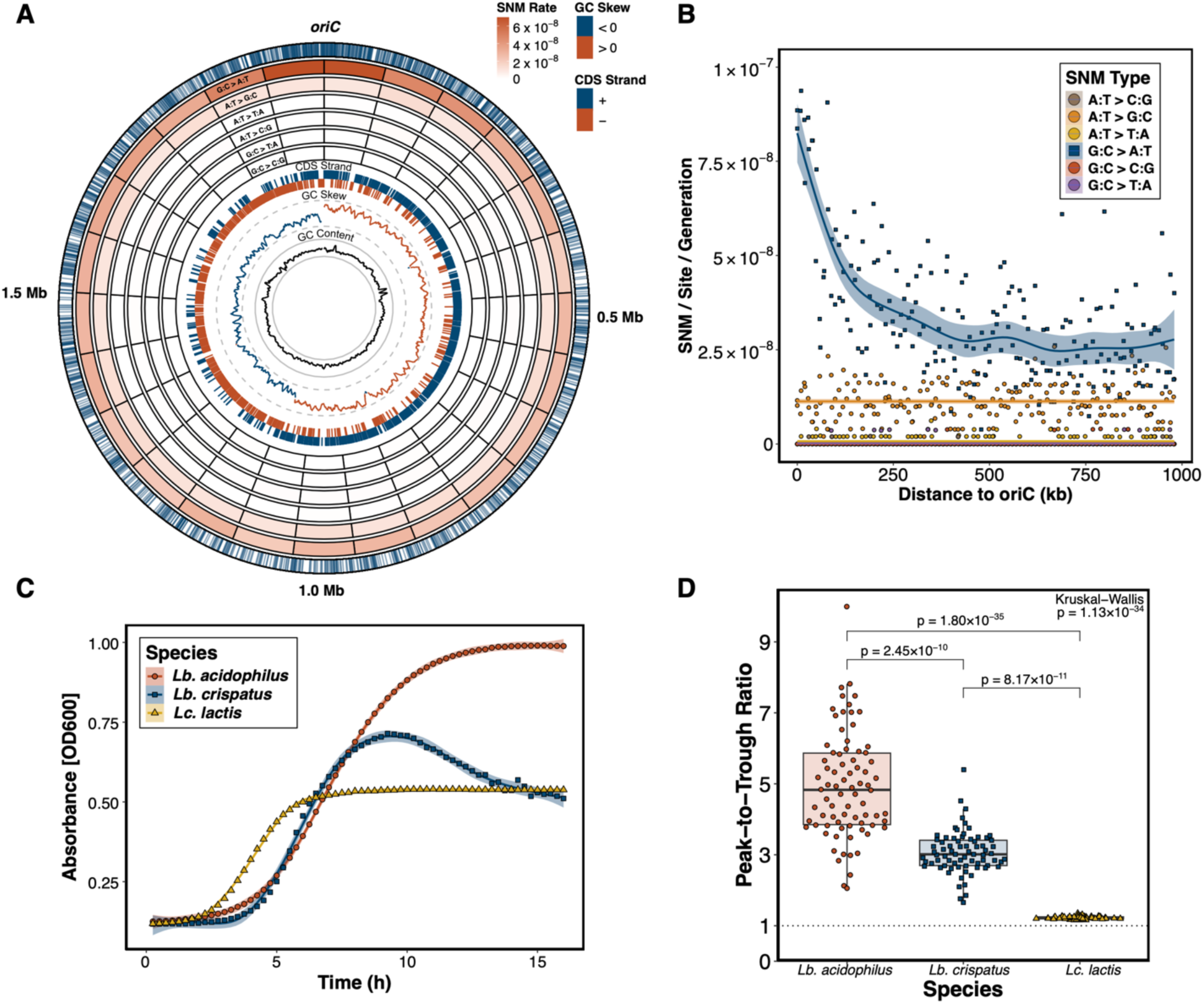
G:C → A:T transitions in *Lb. acidophilus* occur at a higher rate near the *oriC*. (A) Circular genome plot of the *Lb. acidophilus* chromosome. From the outside in: labels indicating genomic coordinates in megabases, blue ticks showing the locations of all observed G:C → A:T transitions, heatmap showing the per site per generation rate of each of the 6 SNM types (from the outside in: G:C → A:T, A:T → G:C, A:T → T:A, A:T → C:G, G:C → T:A, and G:C → C:G) across 25 evenly sized (79154 bp) bins, lines indicating the start positions of each CDS colored by strand, local (1000 bp) GC skew colored by sign with dashed gray scale lines at 0.25 (outer) and −0.25 (inner), and local (1000 bp) GC content with solid gray scale lines at 50% (outer) and 25% (inner). (B) Relationship between distance to the *oriC* and the per site per generation rate of each of the 6 SNM types across 10 kb bins. Square points represent G:C → A:T transitions while circular points represent the other 5 SNM types. (C) Growth curves of the ancestral MA strains in MRS under anaerobic conditions. (D) Peak-to-trough ratios of the derived MA lines for each species. The Kruskal-Wallis p-value is in the top right corner and the pairwise Bonferroni corrected Dunn post hoc p-values are above the brackets.

Initiation of replication requires the unwinding of DNA at the origin into single-stranded DNA - which is more susceptible to cytosine deamination (Frederico et al. 1990). As such, we hypothesized the elevated rates of G:C → A:T substitutions near the origin in *Lb. acidophilus* may result from higher rates of replication initiation. Because of the directional nature of bacterial replication, sequencing coverage is expected to peak at the replication origin and decline along the genome toward the replication terminus. A higher ratio of sequencing coverage at the replication origin relative to the terminus suggests increased replication initiation rates and greater exposure of DNA in a single-stranded state near the origin. To quantify this for each MA line, we used iRep (Brown et al. 2016) to quantify the peak-to-trough ratio of sequencing reads and coverage across the length of the chromosome using the same sequencing data used to call the mutations. We found that the mean peak-to-trough ratio of *Lb. acidophilus* is significantly higher than that of *Lb. crispatus* and *Lc. lactis* (**Fig. 3C**, Kruskal-Wallis p < 10^−33^, Dunn post hoc all p < 10^−9^). To further assess the possibility that *Lb. acidophilus* experiences more time undergoing replication than the other two species, we performed growth curves cultivating the MA ancestor strains for each species in a microplate reader under anaerobic conditions. Our results show that *Lb. acidophilus* spends more time in logarithmic phase than *Lb. crispatus* and *Lc. lactis* (**Fig. 3D**). These results are consistent with the hypothesis that increased replication initiation rates in *Lb. acidophilus* contribute to higher substitution rates at the origin.

### Small indel mutations are also elevated in LAB

In addition to elevated SNM rates, anaerobically cultured LAB had high rates of small (< 50 bp) insertions and deletions (indels). In total, we observed 1042 indels in *Lb. acidophilus*, 71 in *Lb. crispatus*, and 19 in *Lc. lactis*, resulting in respective rates of 6.82 x 10^−9^ (SE: 3.18 x 10^−11^), 4.57 x 10^−10^ (SE: 6.50 x 10^−12^), and 1.48 x 10^−10^ (SE: 5.21 x 10^−12^) indels per site per generation (**Table 1**, **Fig. 4A**). Similar to their SNM rates, these results place LAB on the high end of studied bacterial indel rates but are in line with previous MA/WGS results in *Lc. lactis* (1.14 x 10^−10^) (Long et al. 2018), *S. pneumoniae* (8.9 x 10^−10^) (Jiang et al. 2024), and *S. suis* (2.39 x 10^−10^) (Murray et al. 2021). All three species had higher rates of small deletions than insertions, which is a commonly observed phenomenon across bacteria (Mira et al. 2001).

**Figure 4.**
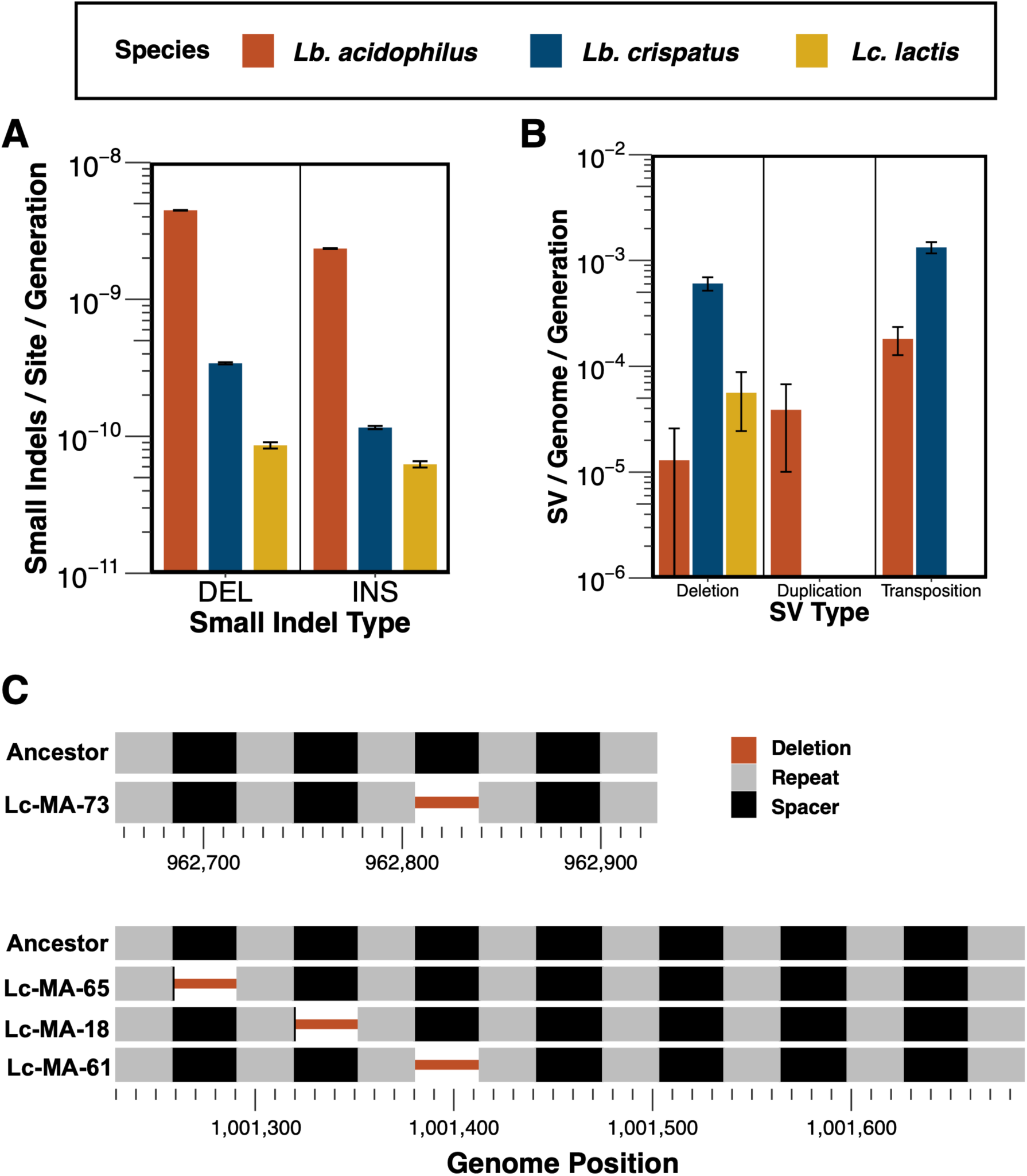
Rates of indels and SVs as well as the locations of deleted CRISPR spacers in *Lb. crispatus*. (A) Per site per generation rates of small (<50 bp) indels. Error bars represent 1 SEM. (B) Per genome per generation rates of SVs. Error bars represent 1 SEM. (C) Locations of CRISPR spacer deletions at the two arrays in *Lb. crispatus*. Thick lines represent intact repeat (gray) and spacer (black) sequences. Thin red lines represent deleted regions.

### IS elements were active in the Lactobacilli but not Lc. lactis

A previous MA/WGS experiment revealed an increased transposition rate of insertion sequence (IS) elements in anaerobically cultured *E. coli* relative to aerobically grown controls (Shewaramani et al. 2017). To determine if anaerobically cultured LAB similarly exhibit elevated IS element transposition rates compared to aerobically cultured bacteria, we profiled structural variants (SVs) with GRASPER (Lee et al. 2014). While our estimates of structural variant mutation rates are conservative due to the limitations of short-read sequencing, we identified 155 SVs that arose during the MA: 18 in *Lb. acidophilus*, 134 in *Lb. crispatus*, and 3 in *Lc. lactis* (**Fig. 4B**). Most SVs observed in the *Lactobacilli* were transpositions caused by IS elements, whereas all 3 SVs found in *Lc. lactis* were deletions. In *Lb. crispatus*, 92 (69%) of SVs were transpositions caused by IS elements, with 87 (95%) caused by IS256 family transposons. The observed rates of IS element transposition in the *Lactobacilli* are comparable to other aerobically cultured bacteria, as rates of 2.10 x 10^−4^ and 2.50 x 10^−3^ transpositions per genome per generation were previously reported for *Burkholderia cenocepacia* and *Deinococcus radiodurans*, respectively (Nzabarushimana and Tang 2018) (**Fig. 4B**).

### CRISPR spacers are deletion hotspots in Lb. crispatus

Clustered, regularly interspaced, short palindromic repeat (CRISPR) arrays are part of a bacterial adaptive immune system (Marraffini 2015). CRISPR loci consist of short repeat sequences that flank spacer sequences commonly derived from infectious mobile elements, such as phage. These spacer sequences serve as a molecular memory and provide the capability to resist infection by parasites that have previously infected an individual’s ancestor. We observed four distinct spacer sequence deletions across the two annotated CRISPR loci in *Lb. crispatus* (**Fig. 4C**). In two cases, the deletion left behind a single base pair of the spacer, while in the other two cases, the deletion removed the entire spacer. While the exact mechanism behind these mutations is unknown, it is unlikely that these deletions are due to misalignment during replication because both flanking repeats were left intact (Garrett 2021).

## Discussion

Lactic acid bacteria (LAB) are foundational members of the human microbiome and are crucial for numerous industrial processes (Bull et al. 2013; Song et al. 2017; Stewart et al. 2018; Shao et al. 2019). Additionally, their facultative to aerotolerant anaerobic physiology makes them ideal candidates for studying how oxygen-tolerant lifestyles affect mutation rates. Here, we report the results of a mutation accumulation experiment performed for over 1,000 generations across at least 50 replicate lines from each of three industrially and biomedically important LAB species. Our study reveals exceptionally high mutation rates and strongly biased spectra in *Lb. acidophilus*, *Lb. crispatus*, and *Lc. lactis* relative to the aerobic bacteria species investigated in previous MA/WGS studies. Moreover, these findings align with results from MA/WGS studies of the aerobically cultured LAB *Lc. lactis* (Long et al. 2018), *S. pneumoniae* (Jiang et al. 2024), *S. suis* (Murray et al. 2021), and the anaerobically cultured archaeon *T. eurythermalis* (Gu et al. 2021), suggesting that reduced oxygen tolerance may be associated with increased mutation rates, even in minimal oxygen environments. As pairwise diversity estimates from MLST data suggest these LAB species represent smaller N_e_ stemming from their domesticated and host-associated lifestyles (Passerini et al. 2010; Xu et al. 2014; Song et al. 2016; Bobay and Ochman 2018; Moyers et al. 2018; Liu et al. 2022; Lynch et al. 2023), our results are consistent with the drift barrier hypothesis of mutation rate evolution (Sung et al. 2012).

One factor possibly contributing to the higher mutation rates observed in LAB species is their characteristic peroxide production attributed to their highly reactive enzymes including pyruvate oxidase, lactate oxidase, NADH oxidase, and NADH-dependent flavin reductase (Condon 1987; Hertzberger et al. 2014; Zotta et al. 2017) and the minimal presence of superoxide scavenging proteins encoded in their genomes (**Fig. 1**). Unlike *E. coli*, a facultative anaerobe that thrives in aerobic conditions and has a mutation rate approximately 90-fold lower than *Lb. acidophilus*, LAB species typically lack catalase (*katE*/*katG*) and superoxide dismutase (*sodA*/*sodC*) genes—two key enzymes responsible for scavenging endogenously produced peroxide and oxygen radicals. We observe strongly biased mutation spectra in LAB, with G:C → A:T transitions representing the majority of SNMs. Although oxidative damage is associated with increased rates of G: C → A:T mutations, prior investigations suggest that LAB produce only trace amounts of H_2_O_2_ under hypoxic and anaerobic conditions (Tomás et al. 2003; Hertzberger et al. 2014). As such, either LAB are sensitive to the minimal oxygen conditions of the anaerobic chamber (∼20ppm of O_2_) or other factors contribute to G:C → A:T bias in anaerobic conditions.

One piece of supporting evidence suggesting that factors outside of oxidative damage can produce a G:C → A:T biased mutation spectrum is the observation of a similar G:C → A:T bias when assessing the anaerobic mutation rates of *E. coli* (Shewaramani et al. 2017). Here, Shewaramani et al. report almost identical G:C → A:T mutation rates in aerobic and anaerobic conditions when cultivating their *E. coli* MA lines on minimal glucose media - which can promote fermentation in aerobic and anaerobic atmospheres. Fermentative metabolism characteristically produces organic acid byproducts and acidic pH can promote DNA damage through protonating cytosine and adenine, increasing the potential for deamination (Wang and Hu 2016). Given that the cultivation of our LAB MA lines is similarly fermentative and that the back mutation of A:T → G:C also occurs at an elevated rate, if not nearly identical in *Lb. acidophilus*, spontaneous deamination may be a major contributor to the enhanced mutation rates of LAB.

Despite the elevation of G:C → A:T and A:T → G:C mutation rates in *Lb. acidophilus* nearly offsetting each other genome-wide and resulting in a relatively balanced AT mutation bias, we observe an unexpected pattern regarding the chromosomal distribution of G:C → A:T transitions. In *Lb. acidophilus*, the G:C → A:T transition rate is three times higher in the 250 kb flanking the replication origin compared to the rest of the genome. Previous studies investigating genetically engineered mutator strains have identified genome position-dependent SNM rate variation (Long et al. 2015; Dettman et al. 2016; Dillon et al. 2018). However, rather than mutation rates peaking in the region flanking the origin, these studies describe a peak in mutation rate mid-chromosome between the replication origin and terminus. Alternatively, in eukaryotes, high mutation rates have been reported near specific human replication origins (Murat et al. 2022). In these cases, multiple SNM types drive the increased mutation rates, with the cause attributed to mechanisms that predominantly affect the immediate vicinity of the origin, such as abortive topoisomerase activity (Murat et al. 2022).

Since G:C → A:T transitions exclusively drive the mutational enrichment in *Lb. acidophilus,* and the enrichment extends significantly further from the origin (∼250 kilobases beyond the *oriC* in *Lb. acidophilus* compared to ∼2.5 kilobases in humans), we propose that this enrichment arises from a distinct underlying mechanism: the unique vulnerability of single-stranded DNA in the prokaryotic replichore (Frederico et al. 1990). This effect is likely amplified in *Lb. acidophilus* due to its higher replication initiation rates and prolonged time spent in the logarithmic growth phase during MA, compared to *Lb. crispatus* and *Lc. lactis*. As more mutation studies are conducted, it is essential to examine the positional enrichment of mutations to determine whether an increased rate of G:C → A:T transitions near the origin is a common feature of microbes that thrive in low-oxygen conditions. If so, further investigation is needed to understand how this localized mutation bias influences the genomic organization of anaerobic prokaryotes.

In contrast to the unique positional mutation bias observed in *Lb. acidophilus*, *Lc. lactis* exhibits an exceptionally strong AT mutation bias. While an AT-biased mutation spectrum is not uncommon in prokaryotes (Hershberg and Petrov 2010), the bias in *Lc. lactis* is unusually pronounced, with an equilibrium GC content of just 8% (**Fig. 2B**), significantly lower than its genomic GC content of 35%. Given this discrepancy, factors beyond mutation must contribute to maintaining the genomic GC content, such as GC-biased gene conversion and selection. GC-biased gene conversion is most commonly discussed in diploid eukaryotes and can maintain genomic GC content by favoring GC alleles when resolving DNA heteroduplexes formed during homologous recombination (Duret and Galtier 2009). However, the low rate of homologous recombination in *Lc. lactis* suggests that the impact, if any, of gene conversion on genome composition is minimal, even if compositionally biased (Torrance et al. 2024). Alternatively, selection can also influence genome composition, with factors such as nitrogen availability (Mcewan et al. 1998), exposure to oxygen (Naya et al. 2002), temperature (Musto et al. 2004), and codon usage (Sueoka 1962) serving as possible sources of selective pressures. The minimum GC content required to encode all 20 amino acids is estimated to be around 20–25% (Sueoka 1962), significantly higher than the 6.4% equilibrium GC content predicted for *Lc. lactis*. While some bacterial genomes have GC contents as low as 15%, these are typically obligate host-associated species (Lightfield et al. 2011; McCutcheon and Moran 2012). This suggests that selective pressure to preserve a functional coding genome likely plays a key role in maintaining *Lc. lactis*’s 35% GC content.

Lastly, Lb. crispatus also exhibits unique mutational characteristics. In addition to its elevated SNM rates, we identified four independent deletions within its CRISPR loci (**Fig. 4C**). These deletions removed most or all of a CRISPR spacer while leaving the surrounding repeats intact. CRISPR activity is particularly significant in LAB, as foundational discoveries describing CRISPR-mediated phage defense were first made in *Streptococcus thermophilus*, a key species in yogurt and cheese production (Barrangou et al. 2007). Our literature search revealed examples where both the spacer and its adjacent tandem repeat were deleted (Gudbergsdottir et al. 2011; Lopez-Sanchez et al. 2012; Citorik et al. 2014; Rao et al. 2017; Stout et al. 2018), but we found no reports of deletions affecting only the spacer. It is possible that losing only the spacer could disrupt CRISPR RNA maturation or interfere with the incorporation of new spacers into the array. Either scenario could weaken the bacterium’s defense against genomic parasites, making such deletions subject to strong negative selection in both wild and industrial strains. Alternatively, our findings suggest that CRISPR locus rearrangements may occur through a multi-step process, beginning with the loss of a spacer, which then increases the likelihood of losing the adjacent repeat. In this context, the *Lb. crispatus* ATCC 33820 strain could serve as a valuable model for studying the mechanisms that maintain CRISPR arrays.

In conclusion, the MA/WGS method has proven to be a powerful tool for characterizing mutation rates and patterns across the tree of life. To fully understand how mutation rates shape and are shaped by evolution, it is essential to consider the environmental contexts in which they arise. Here, by analyzing the high mutation rates of aerotolerant anaerobes under anaerobic conditions, we provide further evidence that the mutation dynamics of anaerobic species differ from those of organisms adapted to atmospheric oxygen levels while still aligning with the drift-barrier hypothesis of mutation rate evolution. We also demonstrate an abundance of mutations associated with spontaneous deamination, highlighting the inherent fragility of purines even under conditions that minimize oxidative stress. Given these high mutation rates, careful handling when maintaining lactic acid bacteria cultures is crucial in laboratory and industrial settings. Our comparisons to aerobically cultured *Lc. lactis* suggest that exposure to oxygen-rich conditions can accelerate mutation rates, potentially leading to unintended consequences for stock populations. Ultimately, our work illustrates the importance of studying mutation rates in fastidious microbes less adapted to aerobic conditions. As aerotolerant and obligate anaerobes represent a significant fraction of the human microbiome and include key species in food and bioenergy production, further investigation into the mutation properties of these taxa is essential.

## Materials & Methods

### Bacterial Strains

We acquired the three strains that served as ancestors for mutation-accumulation: *Lb. acidophilus* ATCC 4356, *Lb. crispatus* ATCC 33820, and *Lc. lactis subsp. lactis* ATCC 11454, directly from the American Type Culture Collection (ATCC). Upon receipt, the strains were immediately cultivated as directed by resuspending the entire lyophilized pellet in 0.5 mL of MRS culture media (Remel R454062) and incubating in 95% Air, 5% CO_2_ at 37°C, until turbid. Cultures were then vortexed, and streaked for isolation on MRS agar (Fisher Scientific, BP9744-5), before cryogenically storing stock cultures in 40% glycerol at −80°C.

### Construction of phylogeny and identification of oxidative stress-associated genes

The reference genomes for our MA ancestor strains (*Lb. acidophilus* ATCC genome ID: 4744cca046d94f76, *Lb. crispatus* NCBI accession: NZ_CP072197.1, and *Lc. lactis* ATCC genome ID: 5635357f23ad4cd3) and for *S. pneumoniae* (NCBI accession: NC_008533.2), *S. suis* (NCBI accessions: PRJNA763404), *E. coli* (NCBI accession: NC_000913.3), and *T. eurythermalis* (NCBI accession: CP008887.1) were annotated with the EggNOG-mapper v2 web tool with default settings (Cantalapiedra et al. 2021). To build the phylogenetic tree, sequences of the *rpoB* gene from each species (*S. suis* MA2 was used as a representative for its species) were aligned with MAFFT v7.505 using the auto option (Katoh and Standley 2013). IQ-TREE v2.2.2.6 was used to select the best model and infer the maximum likelihood phylogeny (Kalyaanamoorthy et al. 2017; Minh et al. 2020). Annotations were then manually profiled for genes previously reviewed as related to oxidative stress in the genus *Lactobacillus* (Zotta et al.

2017).

### MA Culture Conditions

To initiate MA, all three bacterial strains were revived in anaerobic conditions (90% N2, 5% CO2, 5% H2; vinyl anaerobic chamber, Coy Lab) by streaking for isolation on MRS agar.

Plates were incubated anaerobically at 37°C by transferring the inoculated plates into an anaerobic jar (BD, 260622) containing an anaerobic atmosphere-generating pack (Thermo, R681001). Anaerobic jars containing plates were only opened within the anaerobic chamber and sealed before removal from the anaerobic chamber. After 48 h of incubation, we randomly selected 96 isolated colonies from each of the revived ancestor plates and restreaked these colonies for isolation on MRS agar, with each MA line occupying a single quadrant of a 4-quadrant petri dish (VWR, U25384-348). To ensure random selection of colonies and reduce the possibility of selection due to bias in colony picking, each plate quadrant was pre-dotted about 0.5 inches from the plate intersection. Lines were subjected to single-cell bottlenecks every 48 h by selecting the colony closest to the dot and restreaking for isolation over the course of 100 days, resulting in 50 transfers. Every 10 transfers the colony second-closest to the dot was selected as a representative colony and was stored for historical record in 96-well plates containing 40% glycerol at −80°C. After the final transfer, the colony closest to the dot was resuspended in a 96-well plate containing 40% glycerol and stored at −80°C.

### Estimating MA Generations

We used a standard protocol to estimate the number of MA generations for each species (Joseph and Hall 2004; Behringer and Hall 2016). Briefly, we obtained single colonies of the MA ancestor of each species by streaking for isolation on MRS agar and incubating anaerobically for 48 h at 37°C. Plates were imaged and the area of 5 random colonies was quantified using Image J (Schneider et al. 2012). The colonies whose area was quantified were then suspended and plated to determine the respective number of colony-forming units (CFU). This process was repeated every 10 transfers to track the average colony population size throughout the MA experiment. Using the average colony population size in CFU at each of the 5 time points, we calculated the number of generations in each 48-hour incubation period as the average of the number of generations, g, estimated for each time point using the following

equation: 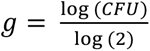.

### DNA Extraction and Sequencing

At the conclusion of the MA experiment, each MA line was struck a final time onto MRS agar and incubated anaerobically for 48 h at 37°C. A loop of cells was collected across the plate to ensure the collection of an appropriate amount of biomass, and that no new novel mutation would have been fixed during the collection of cells for DNA Sequencing. DNA was then extracted with the Qiagen DNeasy UltraClean Microbial Kit (Qiagen, 12224-250), with lysis steps completed under anaerobic conditions. A quality control analysis was performed on the DNA samples using the Qubit HS-dsDNA quantification kit (Qubit, Q32854) to determine quantity before submission to the VANTAGE core for library preparation and DNA sequencing. The samples were normalized to target 100 ng of total input for each sample, and libraries were prepared using the Twist Biosciences kit (Twist, 104207) following a miniaturized version of the manufacturer’s protocol. The libraries were quantitated and pooled for downstream processing. The pool quality is assessed using the Agilent Bioanalyzer and quantified using a qPCR-based method with the KAPA Library Quantification Kit (Kapa, KK4873) and the QuantStudio 12K instrument. Prepared libraries were pooled in equimolar ratios, and the resulting pool was subjected to cluster generation using the NovaSeq 6000 System, following the manufacturer’s protocols. 150 bp paired-end sequencing was performed on the NovaSeq 6000 platform targeting 2M reads per library. Raw sequencing data (FASTQ files) obtained from the NovaSeq 6000 was subjected to quality control analysis, including read quality assessment. Real Time Analysis Software (RTA) and NovaSeq Control Software (NCS) (1.8.0; Illumina) were used for base calling. MultiQC (Ewels et al. 2016) (v1.7; Illumina) was used for data quality assessments.

### Quality Control and Mutation Identification

Raw sequence reads were trimmed and filtered with fastp 0.23.4 (Chen et al. 2018).

Processed reads were then checked for contamination with Kraken2 2.1.3 (Wood et al. 2019). The processed reads from uncontaminated samples were then aligned to their respective reference genomes using BWA 0.7.10-r789 (Li 2013). Alignment metrics were generated with Samtools 1.20 (Danecek et al. 2021). Raw and processed sequence reads were analyzed with FastQC 0.12.1 (https://github.com/s-andrews/FastQC) and these results, along with the Kraken2 output and alignment statistics were visualized with MultiQC 1.19 (Ewels et al. 2016). Duplicate reads in samples without contamination and with sufficient mapped read depth were identified with Picard Tools, and then SNM and small indel variants were identified with the HaplotypeCaller program in GATK 4.1.2.0 (Poplin et al. 2018) with ploidy set to 1. Variant calls from GATK were hard filtered with BCFtools 1.18 (Danecek et al. 2021) using the following filters: F_MISSING = 0, QD ≥ 2, SOR ≤ 5, FS ≤ 60, MQ ≥ 40, MQRankSum ≥ −20, ReadPosRankSum ≥ −8, and QUAL ≥ 30. Only chromosomal variants were considered, so variants called on the plasmids present in *Lc. lactis* were removed. Lines were then manually checked and filtered for cross-contamination. Mutations occurring in close proximity (<50 bp) to one another within the same MA line are likely not caused by independent mutational events, so these mutations were omitted from downstream analyses as has been done previously (Chan and Gordenin 2015; Behringer and Hall 2016). Larger structural variants were identified with GRASPER 0.1.1 (Lee et al. 2014). Structural variants were manually quality-controlled using the Integrative Genomics Viewer 2.15.2 (Robinson et al. 2011). To fully annotate the deletions observed in the CRISPR arrays of *Lc. crispatus*, we used the specialized annotation web tool CRISPRCasFinder 4.2.30 with default options (Couvin et al. 2018).

### Mutation Rate and Spectrum Calculations

For each MA line, the SNM and small indel rates were calculated using the formula *μ* = *m*/*nT*, where μ is the mutation rate per site per generation, m is the number of observed mutations, n is the number of sites analyzed, and T is the number of MA generations for that species. The SNM spectrum was calculated using the same formula with m being the number of observed mutations of a given type and n being the number of sites where that type of mutation could occur. Rates of SVs per genome per generation were calculated for each line using the formula *μ* = *m*/*T*, where m is the number of SVs observed in a line. For each mutation type, the rate for the species was calculated as the mean of the rates for each MA line in that species. Standard errors for mutation rates in each species were calculated using the formula 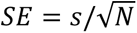, where s is the standard deviation of mutation rates amongst the MA lines and N is the number of MA lines. Mutation rates for each line can be found in **Dataset S1**. Rates for each SNM type in the 10 kb bins analyzed in **Fig. 3** were calculated using the formula *μ* = *m*/*nTl*, where m is the total number of that SNM type observed within the bin for that species, n is the number of sites where that SNM type could occur in that bin, and l is the number of MA lines for that species.

### Mutation Bias Calculations

The AT mutation bias (β) was calculated using the formula *β* = (*GC* → *AT*)/(*AT* → *GC*), where β is equal to the ratio of GC → AT mutations to AT → GC mutations. To calculate fixation bias, we first calculated the equilibrium genome-wide AT composition (ɑ_eq_) that would be expected under neutral evolution from β using the formula *ɑ*_+,_ = *β*/(1 + *β*) (Lynch et al. 2023). When selection acts on genome composition, ɑ_eq_ can be estimated with the formula *α*_+,_ ≈ *βφ*/(1 + *βφ*), where φ is the fixation bias, or ratio of the rate of fixation for new AT alleles relative to that of new GC alleles (Li 1987; Lynch et al. 2016; Lynch et al. 2023). To estimate φ, the formula can be rewritten as *φ* ≈ *ɑ*_*obs*_(*β* − *βɑ*_*obs*_,), where ɑ_obs_ is the observed genome-wide AT composition.

### Effective Population Size Estimates

We used an estimate of N_e_ for *Lc. lactis* calculated from MLST data(Lynch et al. 2023). For *Lb. acidophilus* and *Lb. crispatus*, we estimated the N_e_ using a nucleotide diversity estimate of 0.0042 for *L. delbrueckii* (Song et al. 2016), the type species of the genus *Lactobacillus* that has a similar life history to the two lactobacilli studied here, as it is used in dairy fermentations and is present in the human gut and urogenital microbiomes (Ghosh et al. 2020; Vaughan et al. 2021). To estimate the N_e_ of *Lb. acidophilus* and *Lb. crispatus*, we used the equation *N_e_* = *π*/2*μ*, where π is nucleotide diversity and μ is the per site per generation SNM rate (Tajima 1983).

### External mutation data

To illustrate the relationships between mutational parameters in **Figs. 2B** and **2C**, we used a publicly available collection of mutation data (Lynch et al. 2023). Data for *Mycoplasma mycoides* JCBI-syn1.0 was removed as this is a heavily engineered mutant strain. Data for species with multiple measurements was averaged. To visualize the difference between the SNM rates that were observed and those that would be expected based on the drift-barrier hypothesis for the three anaerobically cultured LAB, we generated a linear regression model using the prokaryotic N_e_ and SNM rate data while excluding the new LAB results. This model was then used to predict the SNM rates of the three anaerobically cultured LAB from their estimated N_e_.

### Examining the relationship between proximity to the oriC and the mutation spectrum

The location of the *oriC* was identified using Ori-Finder 2022 with default options (Dong et al. 2022). The relationship between the distance from the start of the bin to the *oriC* and the rate of each SNM type for each species was tested using Spearman correlation with Bonferroni p-value correction. Peak-to-trough ratios for each MA line were calculated using the bPTR function in iRep 1.1.14 (Brown et al. 2016) with the same BAM files used to call mutations. For the growth curves, the ancestral strains of each species were struck out on MRS agar plates and cultured as described in the MA protocol above. Three colonies per species were inoculated into liquid MRS and incubated overnight shaking at 37°C. From these cultures, 1.5 µL were inoculated into 150 µL of liquid MRS in a 96-well plate. The growth curve was then performed in an Agilent BioTek Epoch 2 plate reader shaking at 37°C, with the optical density at 600 nm measured every 15 minutes.

### Data Availability

All code for data analysis and production of figures can be found on the Behringer Lab’s GitHub at <u>github.com/BehringerLab/LAB_MA_2025</u>. Whole-genome sequencing data generated during this study are publicly available on SRA at NCBI BioProject PRJNA1228167.

## Acknowledgments

We would like to thank B.I.C. and A.J.W. for their helpful comments and suggestions leading up to this study. Anaerobic culture facilities were provided by the Vanderbilt Institute for Infection, Immunology, and Inflammation. Library prep and genomic sequencing were conducted at the Vanderbilt Technologies for Advanced Genomics (VANTAGE). High-performance computing resources were provided by Vanderbilt Advanced Computing Center for Research and Education (ACCRE). This work was supported by National Institute of General Medical Sciences Grant R35GM150625 (M.G.B.), as well as additional funds provided by the Evolutionary Studies Initiative at Vanderbilt (O.F.H. and M.G.B) and the Vanderbilt Undergraduate Summer Research Program (M.Y. and M.G.B.).

---

**Figure S1.**
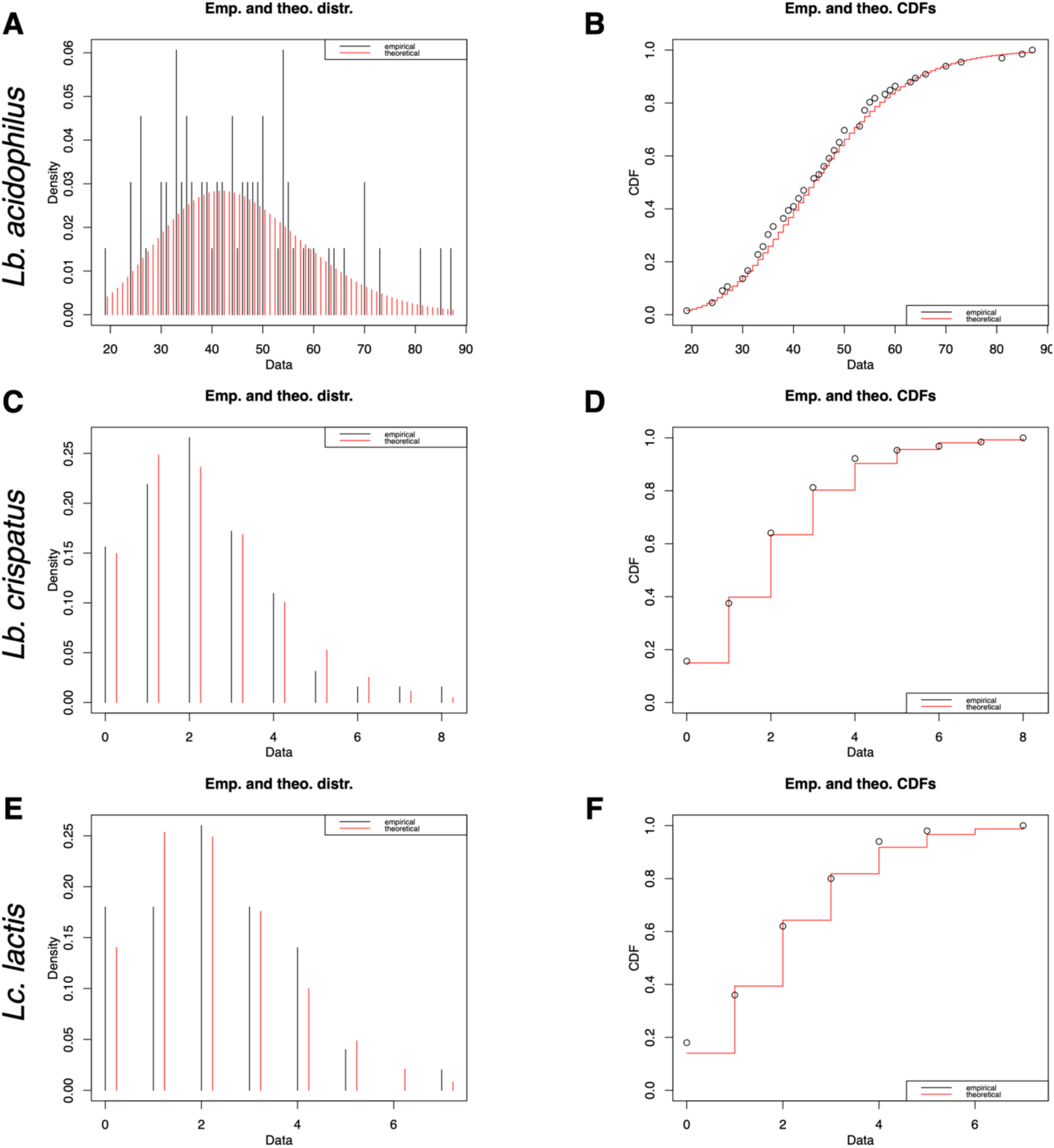
SNM counts fit a negative binomial distribution. Probability density (A, C, and E) and cumulative density (B, D, and F) plots of observed (black) and expected (red) mutation counts given a negative binomial distribution with parameters fit by the R package fitdistrplus.

**Figure S2.**
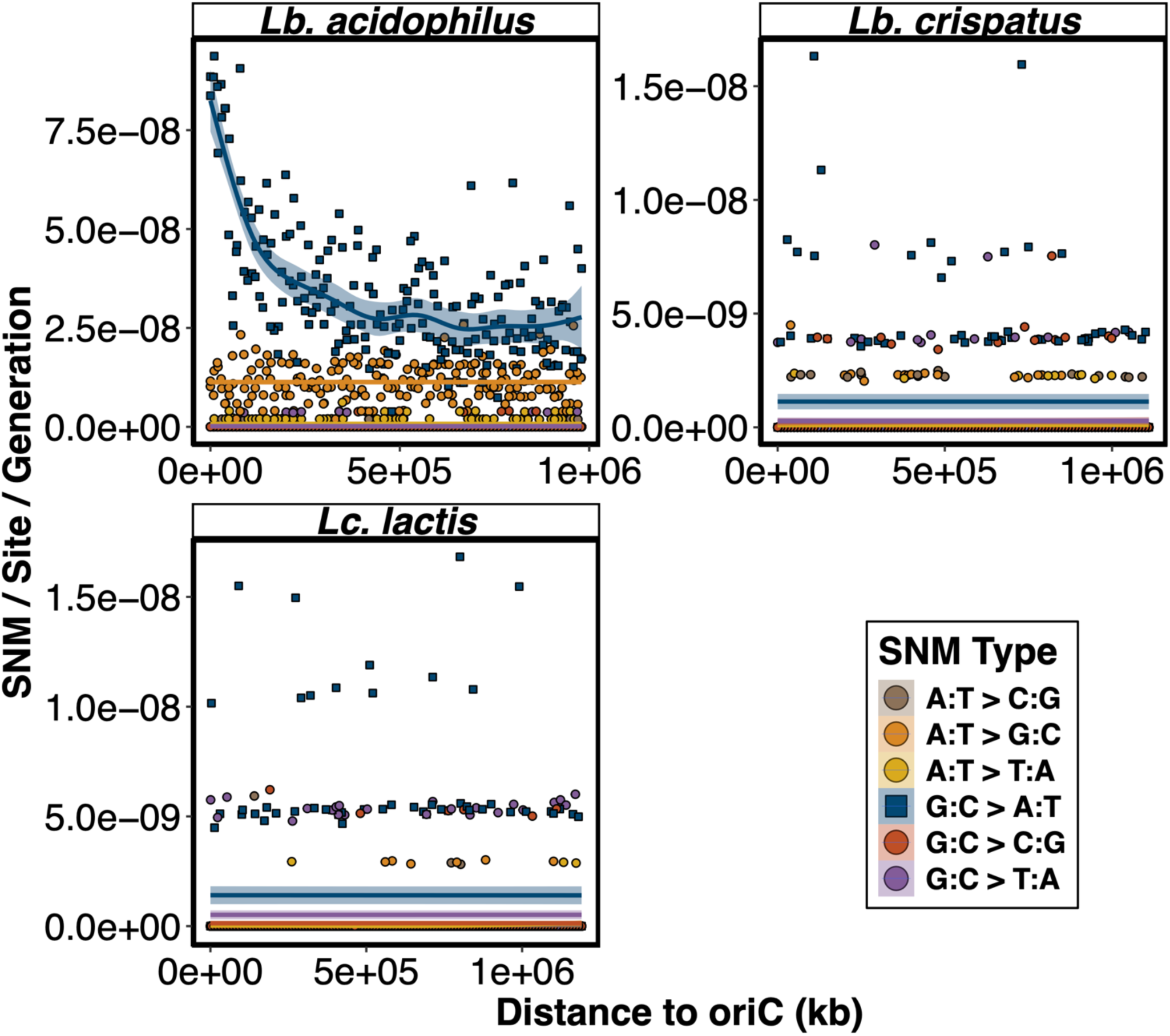
Relationship between distance to the *oriC* and the per site per generation rate of each of the 6 SNM types across 10 kb bins in all three species. Square points represent G:C → A:T transitions while circular points represent the other 5 SNM types.

