## Supplementary figures and images for "Elevated rates and biased spectra of mutations in anaerobically cultured lactic acid bacteria"

### Supplemental Figure 1

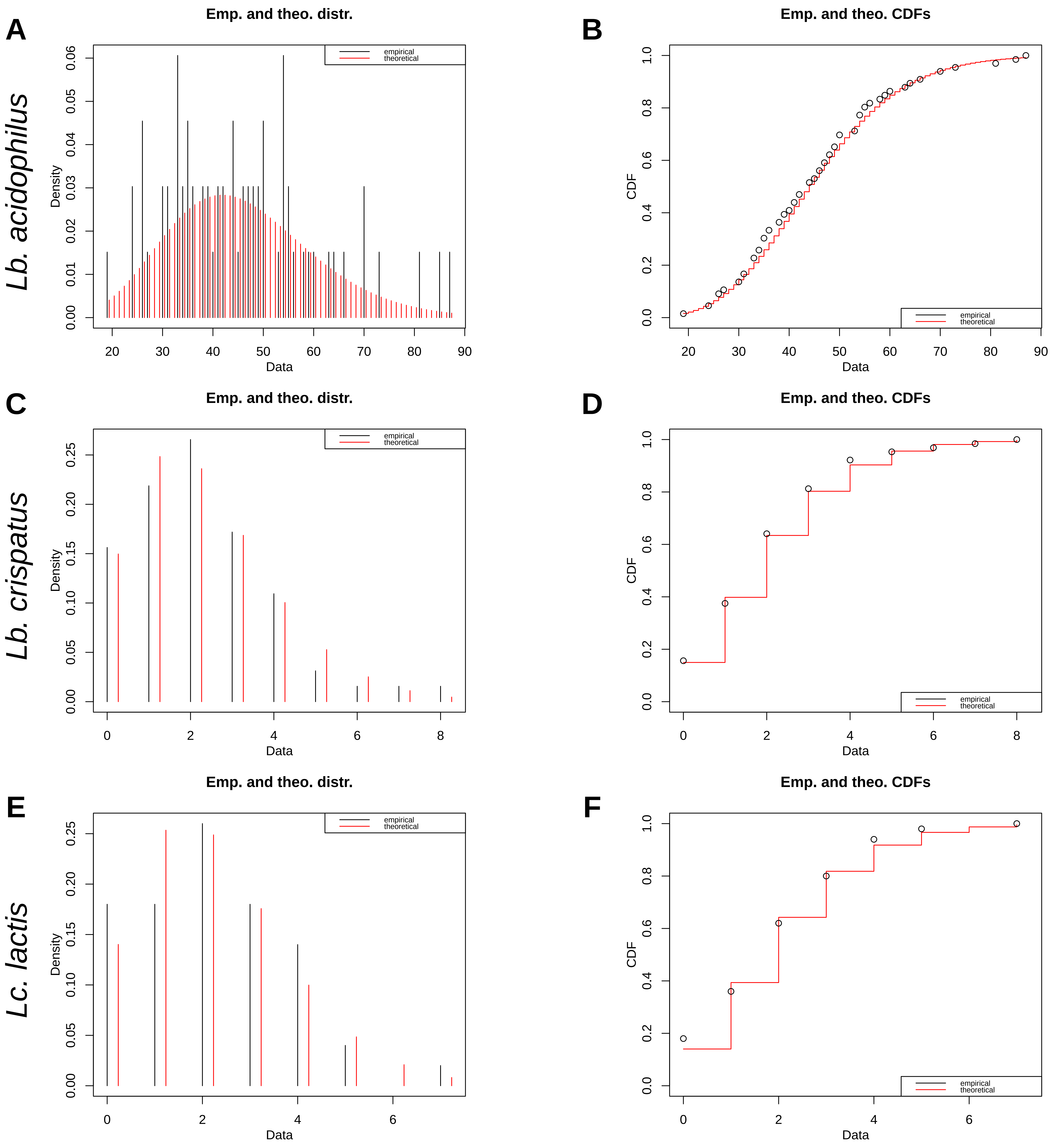

### Supplemental Figure 2

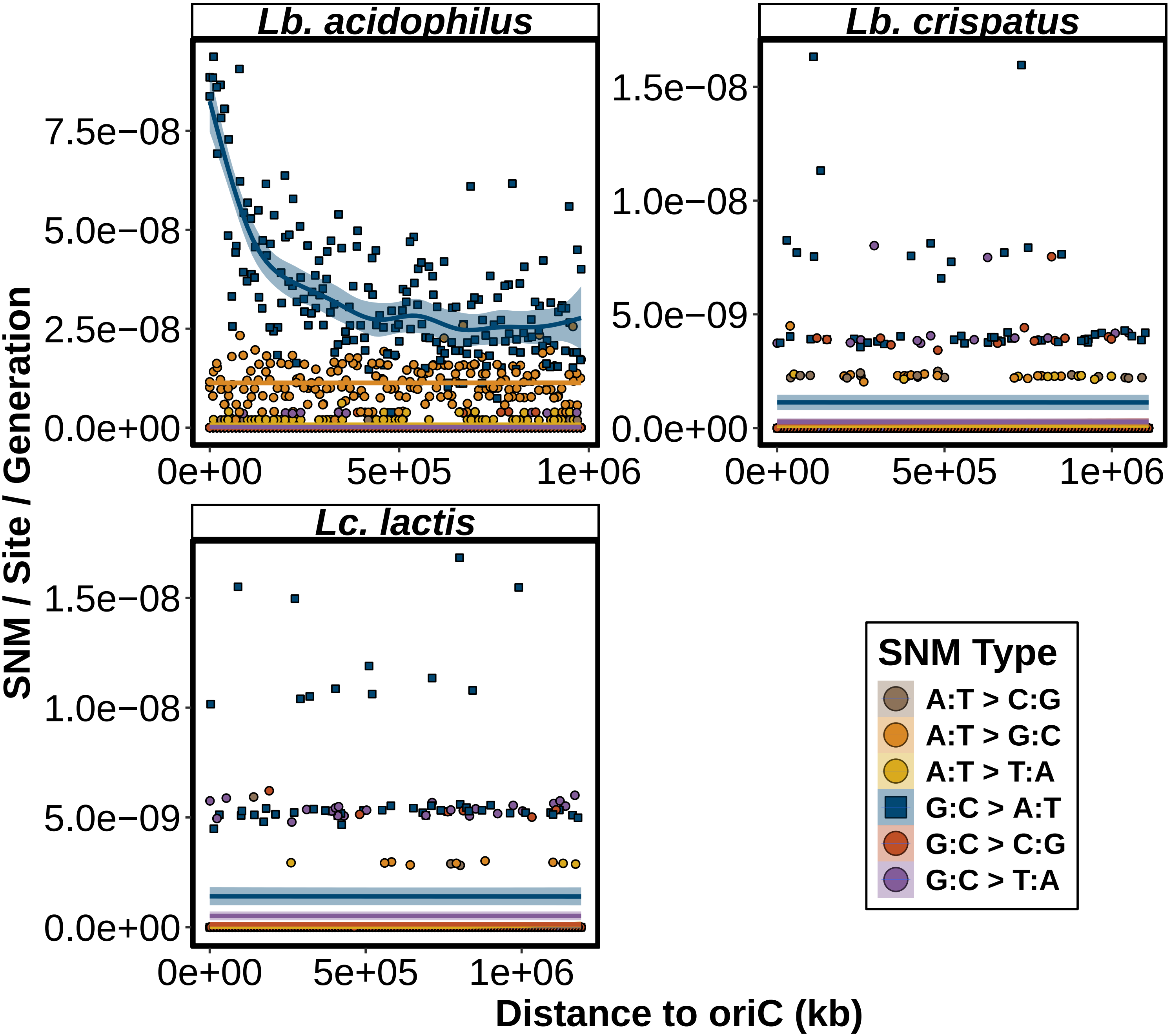
